# CEREBRAL GLUCOSE UTILISATION DURING MUSICAL EMOTIONS: A MULTIMODAL FUNCTIONAL PET/MRI STUDY

**DOI:** 10.1101/2025.08.28.672829

**Authors:** Vesa Putkinen, Andreas Hahn, Jouni Tuisku, Harri Harju, Kerttu Seppälä, Anna K. Kirjavainen, Johan Rajander, Jussi Hirvonen, Lauri Nummenmaa

## Abstract

Functional magnetic resonance imaging (fMRI) studies have demonstrated music-induced activation of the blood-oxygen-level-dependent (BOLD) signal across brain networks associated with auditory perception, motor control, and emotion. However, BOLD-fMRI reflects vascular responses that may not fully capture underlying neural activity. Here, we used simultaneous [^18^F]fluorodeoxyglucose (FDG) functional positron emission tomography (fPET) and fMRI to examine glucose metabolism closely linked to neural activity, alongside hemodynamic responses during pleasurable music listening. Thirty-five female participants listened to self-selected pleasurable music and control stimuli while undergoing 90-minute PET-MRI scans. fPET revealed music-evoked increase in glucose consumption in auditory and motor cortices, as well as reward-related regions, including the nucleus accumbens (NAcc), caudate, insula, and orbitofrontal cortex. The fPET and fMRI results showed substantial overlap though some discrepancies were also observed. Notably, the NAcc exhibited increased glucose consumption in fPET but showed no activation in fMRI. Conversely, deactivation of the default mode network during music processing was only observed with fMRI. These results highlight the complementary nature of neurometabolic and neurovascular processes and offer novel insights into their dynamics during the processing of aesthetic rewards.

## Introduction

Music primes our bodies to move, invites aesthetic engagement, communicates personal and cultural meaning, and, notably, evokes a range of emotions and profound enjoyment. Listening to music places significant processing demands on the brain, engaging auditory processing, motor functions, interoception, memory, attention, as well as various social cognitive and affective functions (Nummenmaa et al., 2021; Vuust et al., 2022; Zatorre & Salimpoor, 2013). Accordingly, functional magnetic resonance imaging (fMRI) studies have shown that music listening increases the blood-oxygen-level-dependent (BOLD) response across auditory, somatosensory, motor, and reward circuits of the brain (Koelsch, 2014; Nummenmaa et al., 2021). BOLD-fMRI, however, measures vascular and oxygenation changes that do not always accurately reflect or originate from neural activity (Drew, 2022). This is highlighted by animal studies showing that neural activity and blood flow can sometimes be decoupled or even inversely related (Huo et al., 2014; Shih et al., 2009). There are also regional differences in the degree of coupling between neural activity and blood flow, which in turn lead to regional differences in how closely the BOLD signal reflects neuronal activity (Shaw et al., 2021). Consequently, music-induced BOLD signal changes may not fully capture the neural activity triggered by music listening. Positron emission tomography (PET) studies have provided complementary evidence by demonstrating the activation of specific neuroreceptor systems during pleasurable music listening (Putkinen et al., 2025; Salimpoor et al., 2011), but such phasic responses can only be measured with the temporal resolution of tens of minutes due to the kinetic properties of commonly used radiotracers.

PET imaging with [^18^F]fluorodeoxyglucose ([^18^F]FDG) provides a measure of brain energy consumption that is closely associated with neural spiking and independent of neurovascular coupling (Sokoloff, 1999) and can therefore reveal metabolic changes not captured by the BOLD signal (Wehrl et al., 2013). Conventional [^18^F]FDG imaging aims at quantifying static, baseline estimates of glucose metabolism (Pak et al., 2022; Rebelos et al., 2021). However, seminal studies have demonstrated that [^18^F]FDG PET can also be applied to measure transient, task-induced changes in glucose metabolism, enabling dynamic quantification with high temporal resolution within a single scanning session (Hahn et al., 2016; Villien et al., 2014) . In this functional PET (fPET) method, the tracer is delivered via a slow infusion so that the slope of the time activity curve dynamically reflects the current cerebral metabolic rate of glucose consumption. This approach provides improved sensitivity to brain-state changes relative to the traditional bolus administration used in static PET scans (Hahn et al., 2016; Villien et al., 2014). Main advantage of this approach is that [^18^F]FDG-fPET provides an absolute, quantified measure of task-specific glucose utilization, unlike the BOLD-fMRI signal that yields a relative and biologically unspecific net index of brain activity (Logothetis, 2008). The feasibility of integrated [^18^F]FDG-fPET and BOLD-fMRI has recently been demonstrated in resting state and cognitive tasks (Godbersen et al., 2023; Hahn et al., 2020) laying the groundwork for investigating the dynamics and coupling between metabolic and neurovascular changes in music processing.

Early fPET studies using simple perceptual and motor tasks suggest a close correspondence between fMRI and fPET signals in sensory and motor regions. However subsequent work with more complex cognitive tasks have revealed divergences between these modalities (Godbersen et al., 2023; Hahn et al., 2020). Recent studies also indicate that fMRI deactivation of the default mode network (DMN) during cognitive tasks is not always accompanied by decreases in glucose consumption (Godbersen et al., 2023; Hahn, Reed, Vraka, et al., 2024; Stiernman et al., 2021), highlighting how [^18^F]FDG-fPET can reveal underlying metabolic activity that is not captured by the BOLD signal. Crucially, the extent to which haemodynamic and metabolic responses converge in brain circuits supporting emotional and aesthetic experiences, such as pleasurable music listening, remains unknown.

### The Current Study

We leveraged simultaneous [^18^F]FDG-fPET and BOLD-fMRI to measure task-evoked changes in cerebral glucose metabolism and haemodynamics while participants listened to self-selected pleasurable music and neutral control stimuli. Current models of the neural basis of music processing, primarily based on fMRI, propose that auditory cortical regions, particularly in the right hemisphere, encode acoustic features and interact with frontal regions to form expectations about the unfolding music (Vuust et al., 2022; Zatorre, 2024). Music also engages motor cortices even in the absence of overt movement, likely to simulate and prepare for moving to the music (Gordon et al., 2018; Zatorre et al., 2007). Music-induced emotions in turn recruit limbic and mesolimbic structures, including the amygdala, hippocampus, and ventral striatum, as well as cortical regions, such as the anterior cingulate, insula, and orbitofrontal cortex, which encode hedonic value and regulate autonomic arousal during emotional experiences across domains (Koelsch, 2014; Mas-Herrero et al., 2021). Given these findings, we expected that pleasurable music would induce heightened BOLD responses in temporal, frontal, parietal and limbic/paralimbic regions and tested whether these responses were accompanied by increases in metabolic activity, indexed by [^18^F]FDG uptake.

## Methods

### Participants

Thirty-five women, recruited through university email lists, participated in the study (mean age ± SD, 24.5 ± 4.7). Only women were included in the study to enhance statistical power, as women tend to report stronger emotional responses across a variety of emotion induction methods (Lench et al., 2011), which could be associated with greater changes in brain glucose metabolism in reward circuits. Screening visiting involved a medical examination, including laboratory blood testing and assessments of general health. The exclusion criteria included a history of neurological or psychiatric disorders, alcohol and substance abuse, current use of medication affecting the central nervous system, and the standard MRI exclusion criteria. Structural brain abnormalities that were clinically relevant or could bias the analyses were excluded by a consultant neuroradiologist. All participants provided written informed consent after receiving a detailed explanation of the study protocol. The study received approval from the Ethics Committee of Turku University Hospital, and the procedures were conducted in accordance with the Declaration of Helsinki.

### Stimuli

Self-selected music was used as stimuli to maximize their pleasurableness (Putkinen et al., 2025; Salimpoor et al., 2011). Before the experiment, the participants were asked to compile a ∼50-minute playlist of music they found pleasurable. We extracted 2-3 45-second representative excerpts from each song to be used as stimuli in the experiment. Most pieces were contemporary pop, R&B, and rap/hip-hop (**See Figure S1**). Control stimuli were 45-second random tone sequences. Tones composing the sequences were 0.1-1 seconds in duration and 110Hz to 1846 Hz in pitch corresponding to musical notes between A2 to G#6 (as in Putkinen et al., 2025).

### Procedure

The participants were asked to fast 5 hours before the scanning. Prior to the scan, the procedure was explained to the participants, their blood glucose concentration was checked to ensure it was below 7 mmol/L, and a urinary pregnancy test was conducted. The participants were instructed to remain still and keep their eyes open throughout the scan. The participants underwent a 90-min simultaneous PET-MRI scanning. The experiment included two 10-min blocks of self-selected pleasurable music and two 10-min blocks neutral control stimulation that were presented in a counterbalanced order across participants. The stimulation blocks were separated by 10-min silent periods (**Figure 1**). To allow sufficient [^18^F]FDG accumulation in the brain, the first block was presented at 15 minutes from the start of the scan. During the stimulation blocks, 45-sec sound stimulation (music or control stimuli depending on the block) was altered with 15-sec silent periods. BOLD data were obtained only during the stimulation blocks. T1-weighted structural brain scans were obtained before the PET scanning.

**Figure 1.**
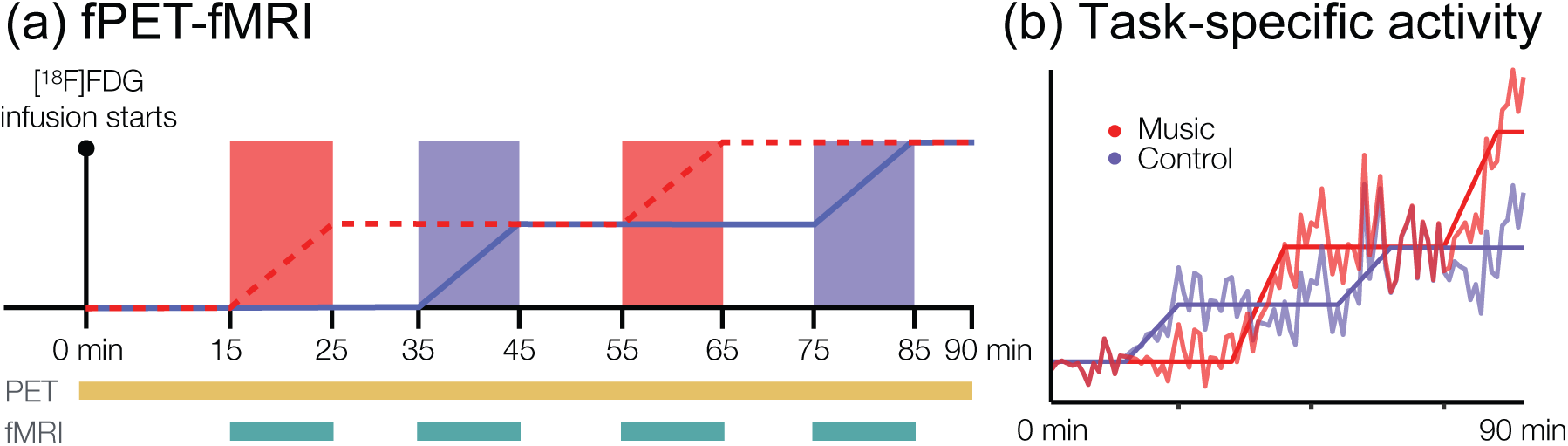
(a) Experimental design. During the 90-min scan, participants listened to self-selected pleasurable music and control stimuli in 10-min blocks. PET data was collected throughout the scan and BOLD data during the stimulation blocks. (b) Task-specific activity in bilateral Heschl’s gyrus from a representative participant. Task-specific activity was obtained by subtracting the contributions of all other regressors (regressor × β) from the raw time–activity curve, leaving only the effect of the task of interest.

### MRI data acquisition and preprocessing

The MRI data were acquired with a 48-channel head coil. High-resolution anatomical T1-weighted (T1w) images were obtained with the MPRAGE sequence for anatomical normalization (1 mm^3^ resolution, TR 8.5 ms, TE 3.7 ms, flip angle 10◦, 256 mm FOV, 256 × 256 reconstruction matrix). For each music and control block, 200 functional volumes were acquired with a T2∗-weighted echo-planar imaging sequence sensitive to the blood-oxygen-level-dependent (BOLD) signal contrast (TR 3000 ms, TE 30 ms, 90◦ flip angle, 256 mm FOV, 128 × 128 reconstruction matrix, 2.7 mm slice thickness, 51 interleaved axial slices acquired in ascending order).

Functional imaging data were preprocessed with FMRIPREP. During preprocessing, each T1w volume was corrected for intensity non-uniformity using N4BiasFieldCorrection (v2.1.0) and skull-stripped using antsBrainExtraction.sh (v2.1.0) using the OASIS template. Brain surfaces were reconstructed using recon-all from FreeSurfer (v6.0.1), and the brain mask estimated previously was refined with a custom variation of the method to reconcile ANTs-derived and FreeSurfer-derived segmentations of the cortical grey matter of Mindboggle. Spatial normalization to the ICBM 152 Nonlinear Asymmetrical template version 2009c was performed through nonlinear registration with the antsRegistration (ANTs v2.1.0), using brain-extracted versions of both T1w volume and template. Brain tissue segmentation of cerebrospinal fluid, white matter and grey matter was performed on the brain-extracted T1w image using FAST (FSL v5.0.9).

### PET data acquisition

PET data were acquired with 3T GE SIGNA PET/MR system (General Electric Medical Systems, Milwaukee, Wisconsin). 271 ± 32 MBq of [^18^F]FDG diluted in saline solution was injected intravenously using a perfusion pump (SpaceStation MRI, B. Braun, Germany). Twenty percent of the tracer was administered as a bolus at the beginning of the scan followed by a constant infusion across the scan duration (Rischka et al., 2018).

### Blood sampling

During the PET scan, manual venous blood samples were drawn at 3, 4, 5, 8, 12, 25, 45, 65, and 85 minutes from the scan onset. Plasma radioactivity was assessed using an automatic gamma-counter (Wizard 1480 3”, Wallac, Turku, Finland).

### PET image processing

The PET data, reconstructed to frames of 1 min, were preprocessed using the automated tool Magia (Karjalainen et al., 2020) (https://github.com/tkkarjal/magia), utilizing preprocessing methods from SPM12 toolbox. MR-based attenuation correction (MRAC) was performed using the zero-echo-time (ZTE) method. Preprocessing included motion-correction of the PET data followed by coregistration of the PET and structural MR images. For the GLM and ROI analyses, pre-processed dynamic PET data were spatially normalized to MNI152 standard space using the deformation fields obtained from the Magia processing.

### GLM

General linear model (GLM) was employed for the modeling of both the fPET and fMRI data. For fPET, the GLM design matrix included regressors for the baseline and the music and control conditions and the intercept, yielding voxel-wise beta-estimates for each regressor. The baseline regressor was defined individually by calculating the mean TAC across all gray matter voxels that were not significantly activated at the group level in the music blocks in the fMRI (cf. Godbersen et al., 2023; Hahn et al., 2020; Rischka et al., 2018). The regressors for the music and control stimulation were defined as linear ramp functions with slope of one during stimulation and slope of zero at other times (Hahn et al., 2016). See Figure 1b for an illustration of auditory cortex activity specific to the music and control conditions in a representative participant (generated using code adapted from Hahn, Reed, Milz, et al., 2024).

The fMRI data were modelled with SPM12 (Wellcome Trust Center for Imaging, London, UK, (http://www.fil.ion.ucl.ac.uk/spm). To reveal regions activated by the music and control stimuli, a general linear model (GLM) was fit to the data where the design matrix included a boxcar regressor for the music vs. silent periods and another boxcar regressor for the control stimulation vs. silent periods.

For both fPET and fMRI, the contrast images for the music and control blocks were then subjected to a second-level analysis for population-level inference in SPM12. Clusters surviving family-wise error rate (FWE) correction (p < 0.05).

### ROI-level cerebral metabolic rate of glucose (CMRGlu)

Blood samples were linearly interpolated to match with PET frame midpoints, creating an input function for modeling. Magia toolbox (Karjalainen et al., 2020) functions were used to calculate Patlak Ki maps from voxel-level 4D data which were obtained by multiplying beta maps with the respective regressors (Hahn et al., 2016). Ki maps were converted to cerebral metabolic rate of glucose (CMRGlu) using the formula CMRGlu = (100*Ki.*Glu)/LC, where LC = 0.676 is the lumped constant in brain. Finally, regional CMRGlu averages were calculated using the AAL atlas (Tzourio-Mazoyer et al., 2002) for regions associated with music processing and default mode network, and Hammer’s atlas (Hammers et al., 2003) for nucleus accumbens (NAc).

## 3. RESULTS

### 3.1. Whole-brain analysis of fPET and fMRI

For the music condition, increased glucose consumption and BOLD responses were observed in the auditory cortex, precentral gyrus, premotor cortex, SMA, insula, and inferior frontal cortex (IFG) (**Figure 2**). Activation in the medial frontal and orbitofrontal cortex, posterior cingulate, and the striatum was only observed in fPET. In contrast, activation of the thalamus, hippocampus, brainstem, and cerebellum was only evident in the fMRI. The music > control contrast revealed similar results with the exception that SMA activation did not survive the cluster correction in the fPET data (**Figure 3**). Compared to the music > baseline contrast, however, this contrast showed weaker effects overall in fPET, especially in the frontal cortex.

**Figure 2.**
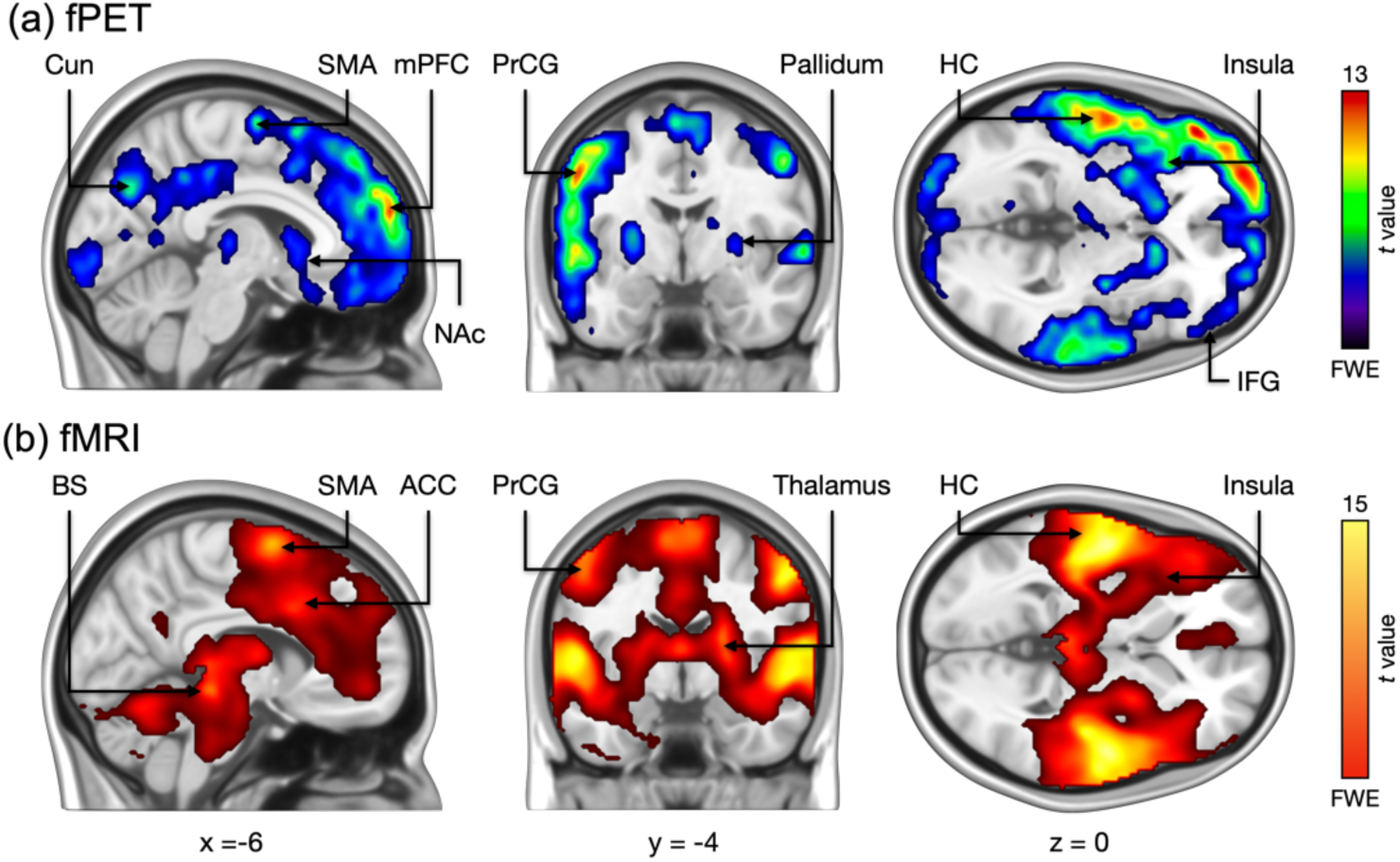
Regions showing higher glucose metabolism (a) and BOLD responses (b) in the music condition versus baseline. ACC = Anterior Cingulate Cortex, BS = Brainstem, Cun = Cuneus, HC = Heschl’s Gyrus, IFG = Inferior Frontal Gyrus, mPFC = Medial Prefrontal Cortex, PrCG = Precentral Gyrus, SMA = Supplementary Motor Area.

**Figure 3.**
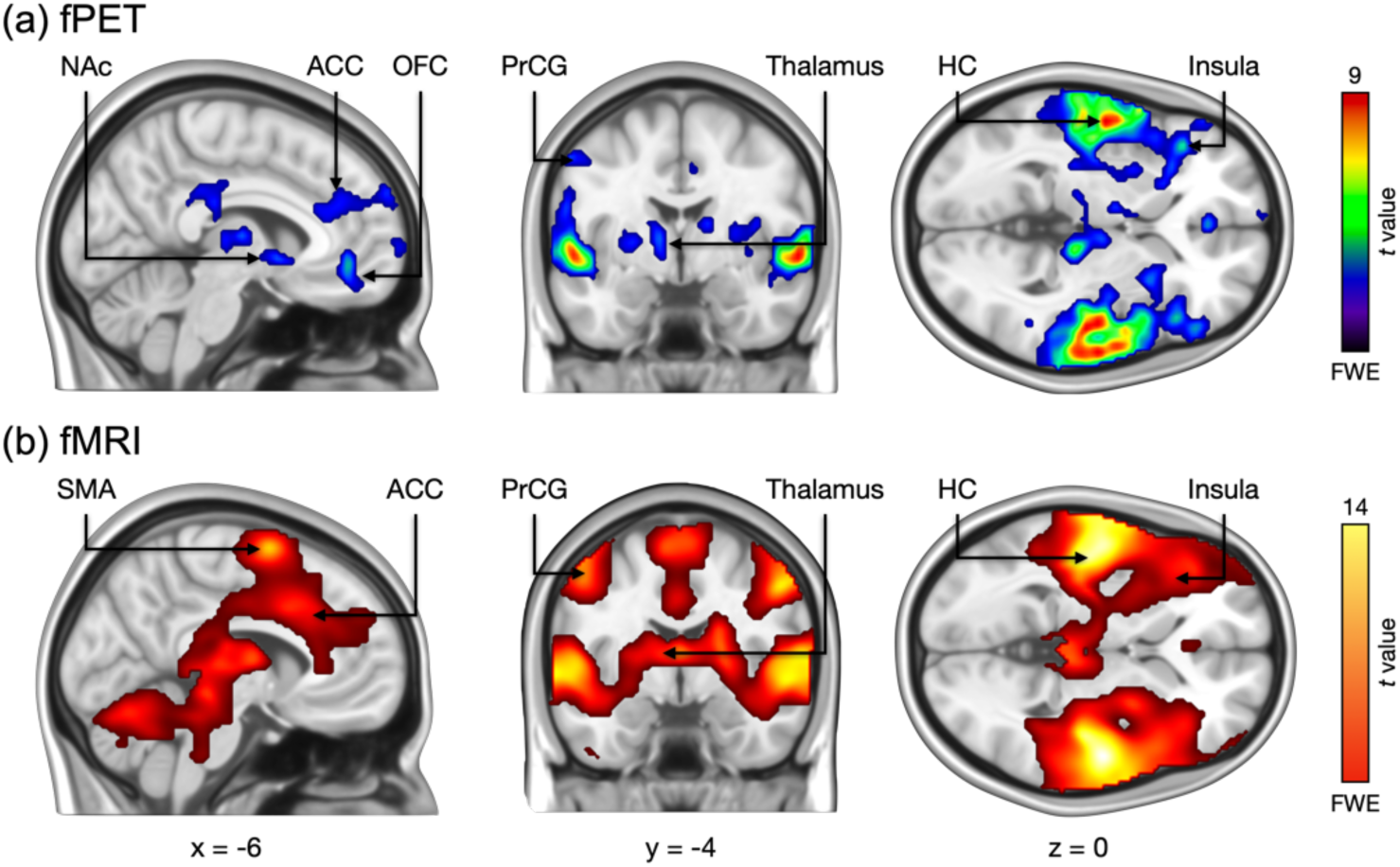
Regions showing higher glucose metabolism (a) and BOLD responses (b) in the music condition relative to the control condition. ACC = Anterior Cingulate Cortex, HC = Heschl’s Gyrus, OFC = Orbitofrontal Cortex, PrCG = Precentral Gyrus, SMA = Supplementary Motor Area.

Overall the fPET and fMRI results showed substantial overlap. The correlation between the unthresholded fPET and fMRI 2^nd^ level T-value maps were .24 and .54 for music > baseline and music > control contrasts, respectively.

Figure 4 shows region where glucose metabolism and BOLD responses were decreased during the music condition relative to baseline. In fPET, strongest deactivations were seen in the cerebellum, brainstem and limbic regions like the amygdala and hippocampus (Figure 4a). In fMRI, deactivations were observed in parietal cortex in the angular gyrus and precuneus as well as in the medial and superior prefrontal cortex (Figure 4b). See the supplementary material for results of the baseline > control and control > music contrasts (**Figure S1**), and for unthresholded effect size maps of the music > baseline and music > control contrasts (**Figure S2**).

**Figure 4.**
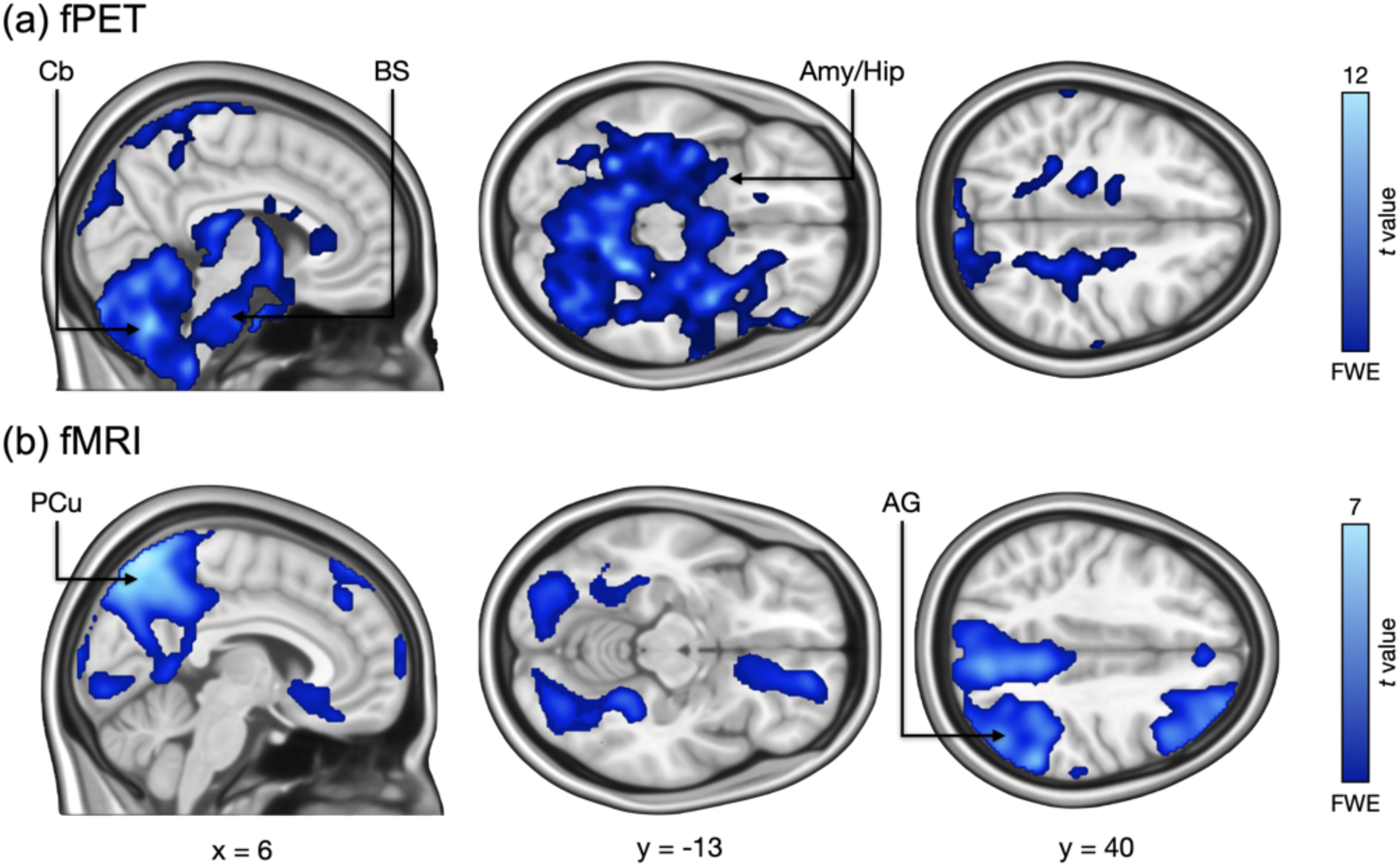
Regions showing deactivations (Baseline > Music) in (a) fPET and (b) fMRI. CB = Cerebellum, BS = Brainstem, Amy/Hip = Amygdala/Hippocampus, PCu = Precuneus, AG = Angular Gyrus.

### 3.2. ROI-level glucose metabolism changes

In the ROI analysis, the largest change in CMRGlu (4.47 μmol/100 g/min) was seen in the left primary auditory cortex in the music condition when compared to the baseline (Figure 5a and Figure 6). In both the music and control conditions, the right auditory cortical regions also showed prominent CMRGlu increases, though these were smaller than those observed in the left auditory cortical regions (t(24) = 4.47, *p* < .001). Conversely, largest decreases in CMRGlu were observed in the amygdala, hippocampus, brainstem and cerebellum for both the music and control conditions. Similar decreases were not seen in the BOLD data, except for regions of the default mode network (Figure 5b). Larger glucose metabolism in the music condition relative to the control condition were seen in the auditory cortices, and thalamus (**Table 1**). The right insula showed smaller decrease in CMRGlu relative to the baseline in the music than in the control condition, resulting in a significant difference between the conditions for the music > control contrast.

**Figure 5.**
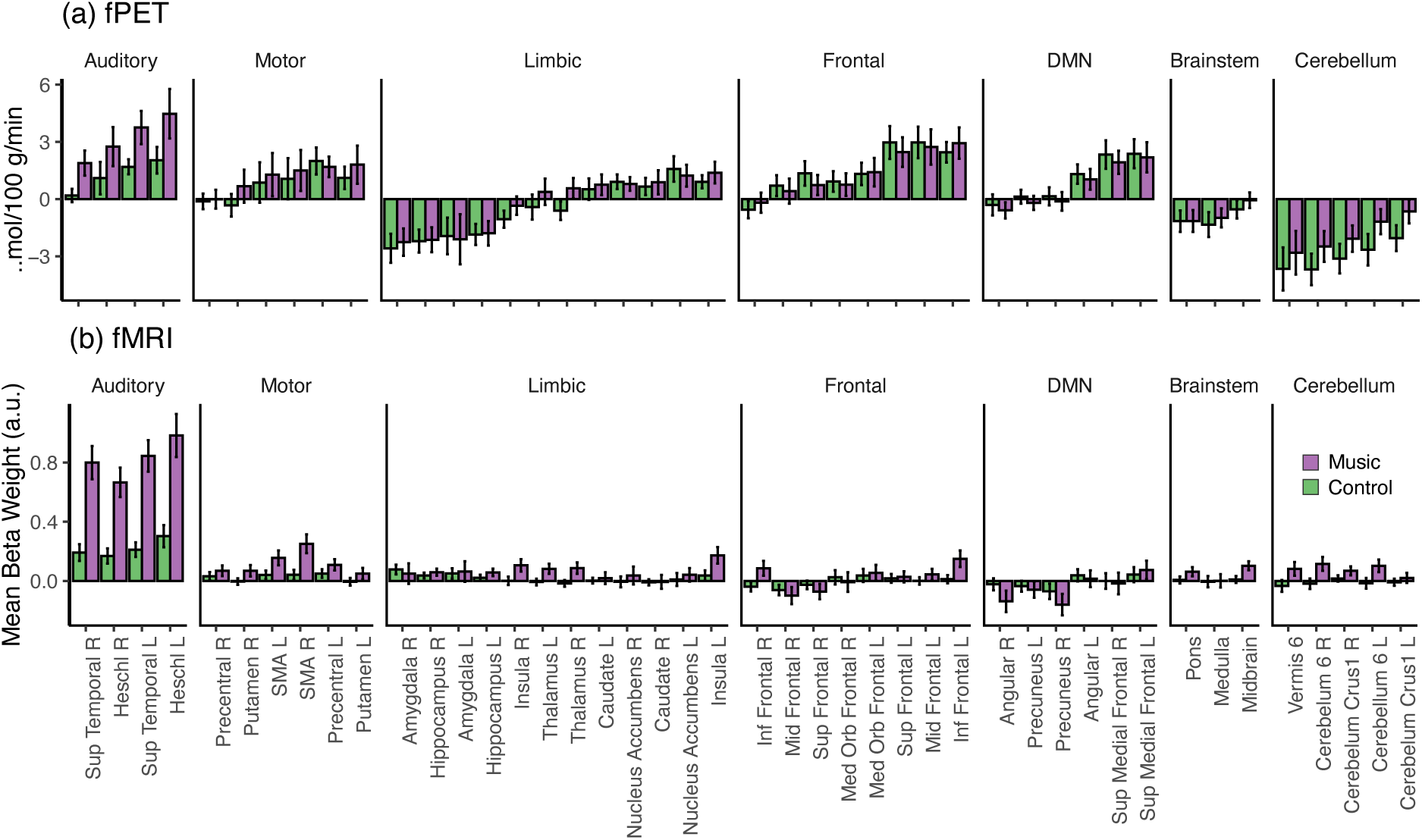
ROI-wise mean CMRGlu for the Music and Control conditions as compared to baseline (a) and fMRI beta estimates (b) for the Music and Control conditions.

**Figure 6.**
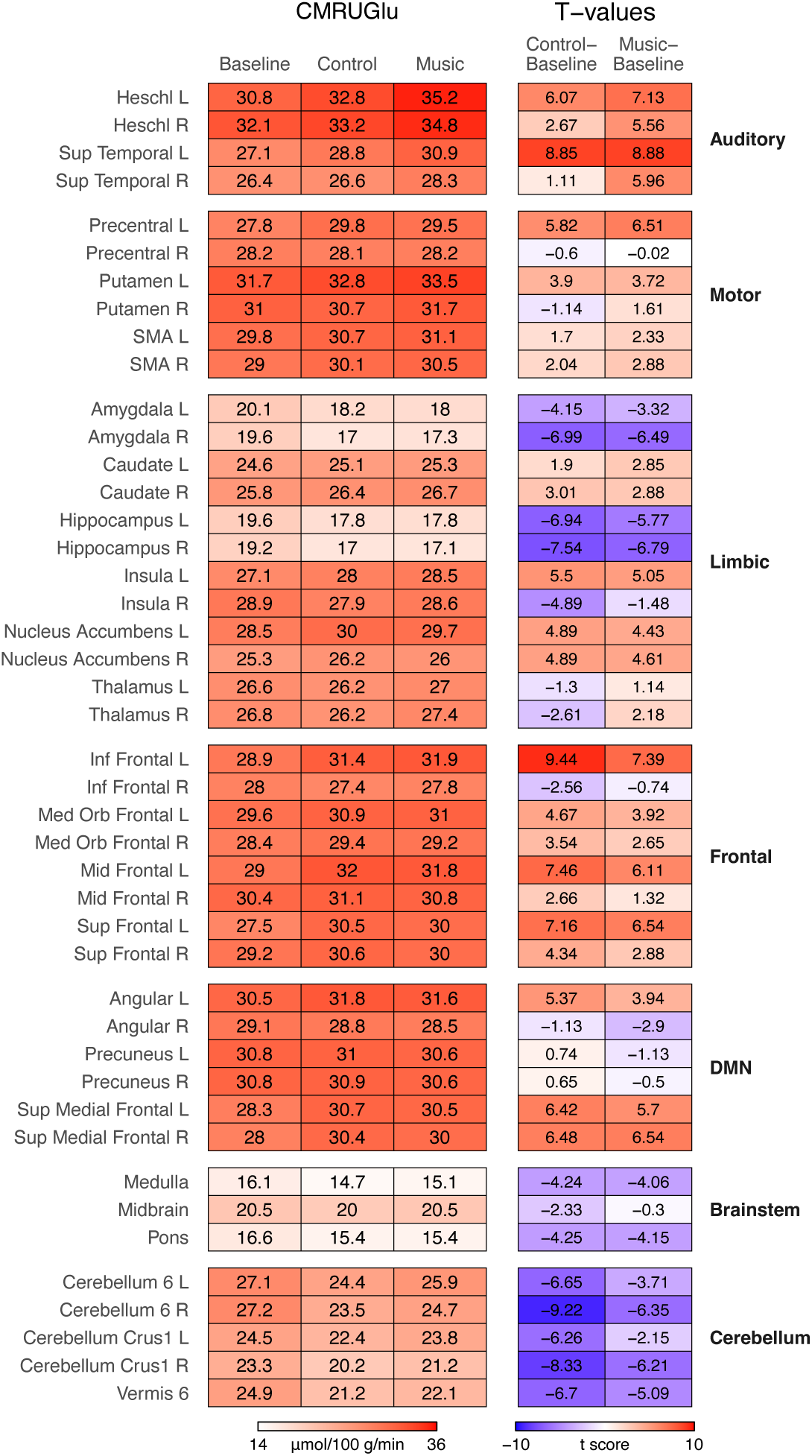
Mean CMRGlu (μmol/100 g/min) for select ROIs for the music, control and baseline and the t-values for the music >baseline, and music >baseline contrasts.

## Discussion

Our main finding was that simultaneous functional [^18^F]FDG-fPET identified increased metabolic responses in auditory, sensory-motor, frontal, and limbic regions during pleasurable music listening. These were paralleled by heightened haemodynamic activity in the same set of regions, yet some notable dissociations between the modalities were also observed. While the nucleus accumbens exhibited increased glucose consumption in fPET, no activation was detected with fMRI. Additionally, key nodes of the default mode (angular gyrus, precuneus, medial prefronal cortex) network showed fMRI deactivation but stable or increased glucose metabolism in fPET. In contrast, the amygdala, hippocampus, brainstem and cerebellum exhibited a marked reductions in glucose metabolism that were not reflected in the fMRI data. These findings provide novel insights into the brain’s energy demands during pleasurable music listening, highlighting both overlapping and distinct metabolic and vascular dynamics underlying sensory, motor, emotional, and cognitive processing of music.

### Music-induced metabolic and haemodymanic changes in the reward circuits

Pleasurable music-listening increased glucose consumption in regions centrally involved in reward processing such as the NAcc, caudate, and orbitofrontal cortex consistent with their domain-general role in reward processing, motivation, and the experience of pleasure (Berridge & Kringelbach, 2015). This first-ever fPET study on central glucose metabolism during music and musical emotions significantly extends previous neuroreceptor PET studies linking striatal dopamine and opioid release to music-induced pleasurable chills (Putkinen et al., 2024; Salimpoor et al., 2011), and fMRI studies reporting music-induced activity in the brain’s reward circuits (Menon & Levitin, 2005; Salimpoor et al., 2013; Trost et al., 2012). By providing direct evidence of increased glucose consumption in reward-related neural circuitry during pleasurable music listening, our findings shed light on the metabolic processes underlying aesthetic reward processing.

In contrast to the fPET results, the fMRI data did not reveal activity in the ventral striatum during the music condition, despite the use of highly pleasurable self-selected music as stimuli(cf. Salimpoor et al., 2011). This aligns with an ALE meta-analysis that found no significant NAcc cluster for pleasant versus unpleasant music, indicating variability in NAcc responses to pleasurable music across studies (Fuentes-Sánchez et al., 2025). The lack of significant striatal activation in fMRI, despite increased glucose utilization, may stem from the differing temporal dynamics heamodynamic and metabolic changes captured by fMRI and fPET, respectively. Hemodynamic responses in the NAcc might be most pronounced during brief moments of music-induced chills or musical uncertainty and surprise—factors not explicitly modeled in this study (Cheung et al., 2019; Gold et al., 2019; Salimpoor et al., 2011). In contrast, fPET effectively integrates metabolic activity over the duration of the entire task block, allowing transient increases in glucose consumption—such as those occurring during moments of music-induced chills—to contribute to the overall signal by increasing the slope of the TAC during the music blocks. Overall, our findings point to the engagement of the ventral striatum in pleasurable music listening, as reflected in increased metabolic demand, even in the absence of concurrent BOLD activation.

We further found that music listening increased glucose consumption and BOLD responses in the insula and cingulate cortex relative to the control condition. Studies linking autonomic arousal indices to BOLD data have implicated these regions in the central regulation of autonomic activity (Ferraro et al., 2022). Music-induced pleasure is strongly associated with autonomic arousal (Salimpoor et al., 2009), suggesting arousal as a potential explanation for the observed insula and cingulate activations. The insula, in particular, is involved in the conscious awareness of bodily states such as heart rate and respiration (Craig, 2002), suggesting that its activity during music listening may reflect the awareness of bodily sensations associated with music-induced emotions (Putkinen et al., 2023).

Haemodynamic activity in the amygdala has been reported in response to both pleasant and unpleasant music (for a meta-analysis, see Koelsch, 2014), aligning with evidence that the amygdala responds to non-aversive yet salient stimuli and broadly contributes to detecting emotional significance (Sander et al., 2003; Pessoa, 2010). Despite this, fPET data revealed a prominent *decrease* in glucose consumption in the amygdala and hippocampus during both the music and control conditions, with no difference between the two conditions (Figure 3a **and S1c**) suggesting higher glucose consumption in these regions at rest than during auditory stimulation. While we observed BOLD responses in the amygdala for the music condition relative to silence, fMRI data showed no significant difference between the music and control conditions in this region. These findings do not support a preferential involvement of the amygdala in pleasurable music processing but instead point to a more general responsiveness to auditory stimulation.

### Music increases glucose consumption in auditory, motor and frontal regions

Listening to pleasurable music increased glucose consumption in the superior temporal gyrus (STG), motor regions (precentral gyrus, premotor cortex and SMA), and the inferior frontal gyrus (IFG). These metabolic changes may reflect the energy demands of forming musical expectations to guide movement (Vuust et al., 2022). Predictive coding models of music processing propose that the STG analyzes musical features such as melody and rhythm, while the IFG maintains this information in working memory, facilitating pattern recognition and expectation generation based on implicit musical knowledge (Zatorre & Salimpoor, 2013). The auditory and frontal regions have been proposed to interact with the motor systems to guide synchronized movement and further aid predictions about the timing of events in the music (Vuust et al., 2022).

As expected, both modalities revealed prominent auditory cortical responses, with fMRI showing particularly strong activation in the auditory cortex compared to other activated regions (Figure 5). Prior studies have found right-lateralized auditory cortical BOLD responses to musical sounds, attributed to greater pitch-processing precision in the right auditory cortex compared to the left. In contrast, we observed higher glucose consumption and stronger BOLD signals in the left auditory cortex. Notably, this leftward lateralization cannot be explained by the presence of lyrics, as the same asymmetry in the superior temporal gyrus (STG) was observed even for control stimuli lacking linguistic content. Evidence supporting rightward lateralization in music processing comes primarily from studies using stimuli specifically designed to disentangle tonal from temporal or linguistic processing, which show left-hemisphere dominance (Zatorre et al., 2002). In contrast, we used naturalistic musical stimuli, primarily contemporary pop songs with strong rhythmic and temporal elements, which may explain the stronger left auditory cortical activation. Since we could not formally analyze how musical features influence [^18^F]FDG uptake in the left versus right auditory cortex, this hypothesis remains to be tested in future studies.

In fMRI, we observed activation in the precentral gyrus, premotor cortex and SMA in the absence of movement, possibly reflecting covert motor simulation, action planning, or the sensation of beat (Matthews et al., 2020). Both imaging modalities also revealed activation in the putamen, a key node of the basal ganglia motor circuit (Lanciego et al., 2012). Cortically, increased glucose consumption was also detected in the premotor cortex, though the difference between control and music blocks was less pronounced than in fMRI and confined to the left hemisphere (**Figures 2 and 3**). Additionally, the cerebellum, which plays a key role in motor control, showed activation in fMRI but exhibited reduced [^18^F]FDG uptake during music listening compared to baseline in fPET. These discrepancies between the imaging modalities suggest that motor preparation or simulation may trigger and increase in blood flow that is disproportionate to the metabolic demands, leading to partial uncoupling between haemodynamics and glucose consumption in the motor system.

### Metabolically costly music-induced DMN deactivation

BOLD responses in regions of the default mode network (DMN), including the angular gyrus and precuneus, were reduced during the music blocks compared to the silent fixation periods between blocks. However, a similar decrease was not observed in glucose consumption during these stimulation blocks in the fPET data. In contrast, fPET revealed increased glucose metabolism during both the music and control blocks relative to the baseline (no music) periods, particularly in the angular gyrus and medial prefrontal cortex—key regions of the DMN. This dissociation between hemodynamic activity and glucose metabolism in the DMN aligns with previous findings comparing cognitive tasks to resting-state conditions. Together, these results further support the notion that task-induced deactivation of the DMN may, paradoxically, be metabolically demanding and require increased energy consumption (Godbersen et al., 2023; Stiernman et al., 2021). Such dissociation has been postulated particularly for tasks with an internal focus and may depend on the cortical network engaged during task (Godbersen et al., 2023; Hahn, Reed, Vraka, et al., 2024).

### Limitations

We scanned only female participants, which may restrict the generalizability of our findings to males. Furthermore, the limited temporal resolution of our fPET measurements (one-minute frames) also poses a constraint for linking dynamics of glucose metabolism and heamodynamic responses. While this resolution allowed us to capture differences in glucose consumption between the music and control conditions, it may have obscured finer, transient changes in metabolism. Future studies employing higher temporal resolution fPET, could provide more detailed insights into the temporal dynamics of glucose consumption and enable direct comparisons of the time courses of metabolic and hemodynamic signals (cf. Hahn et al., 2024). This could be facilitated by the use of modern long axial field of view total-body tomographs which provide significantly improved counting statistics and signal-to-noise-ratio, thus allowing high temporal precision while modelling data with frame length within the rank of seconds (Knuuti et al., 2023; Zhang et al., 2020).

### Conclusions

We conclude that changes in energy metabolism within limbic, auditory, sensorimotor, and frontal regions support musical and aesthetic emotional processing. This study is the first to combine functional [^18^F]FDG-fPET and fMRI to investigate hemodynamic and metabolic activity during the processing of aesthetic rewards. Our findings reveal both converging and dissociable patterns of vascular and metabolic activity underlying the sensory, motor, emotional, and cognitive components of pleasurable music listening.

## Acknowledgments

We sincerely thank Eveliina Rantakylä for her contributions to participant recruitment, data collection and data management and preprocessing.

## Supplementary Materials for

**Figure S1.**
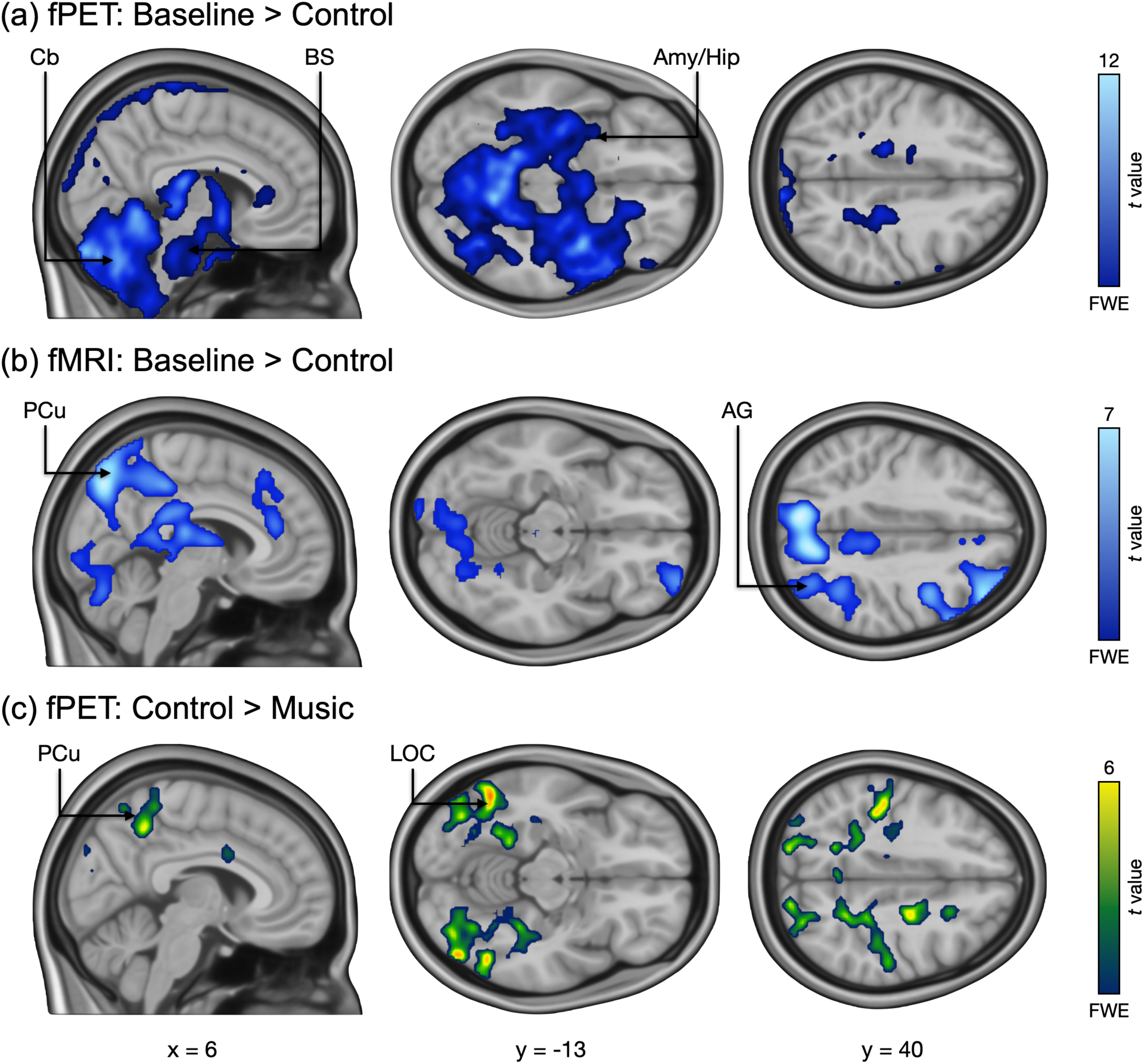
Regions showing deactivations (Baseline > Control) in (a) fPET and (b) fMRI. AG = Angular Gyrus, Amy/Hip = Amygdala/Hippocampus, BS = Brainstem, CB = Cerebellum, LOC = lateral occipital cortex, PCu = Precuneus.

**Figure S2.**
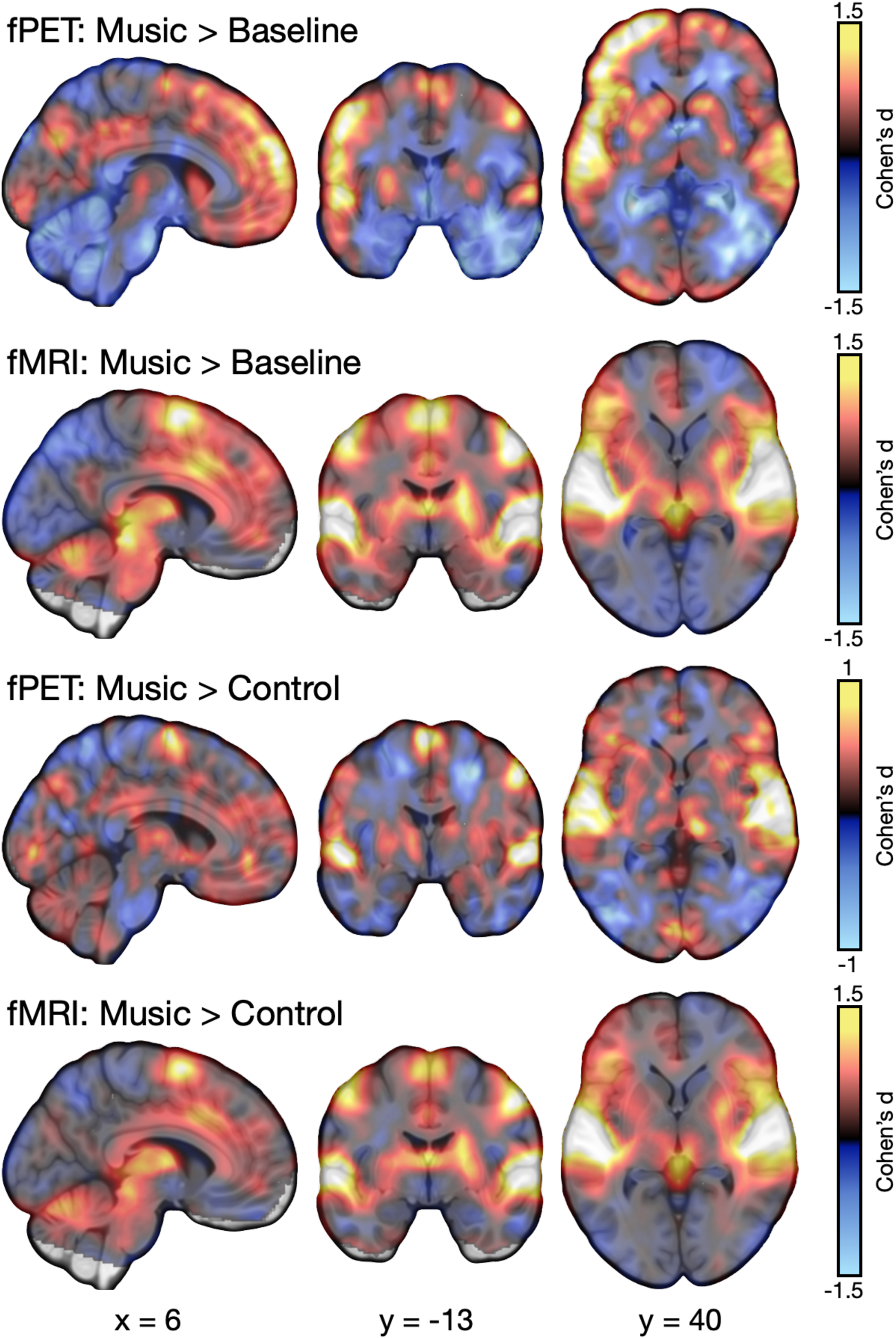
Unthresholded effect size maps for music > baseline and music > control contrast for fPET and fMRI.

